# How the naked mole-rat resists senescence: a constraints-based theory

**DOI:** 10.1101/2021.09.22.461400

**Authors:** Felipe A. Veloso

## Abstract

Here, I present a theory describing how the stabilization of constraints imposed on chromatin dynamics by the naked mole-rat’s histone H1.0 protein—which in terminally differentiated cells constrains the accessibility of the nucleosome core particle for histone-modifying enzymes and chromatin remodeling factors—explains its resistance to both senescence and cancer. Further, this theory predicts that a mutant house mouse displaying such stabilization will be similarly resistant to both senescence and cancer. A proof-of-concept computational analysis is presented and two predictions for the direct testing of the theory are provided. These experiments comprise, as test subjects, mutant naked mole-rats synthesizing a house mouse (Mus *musculus*)-like histone H1.0, and mutant house mice synthesizing a naked mole-rat-like histone H1.0. The predictions are that the constraints on chromatin dynamics embodied by the respective mutant histone H1.0 proteins will negate the otherwise significant resistance to both senescence and cancer of the naked mole-rats and, conversely, confer such resistance to the house mice. A verification of these predictions will imply that constraints on chromatin dynamics embodied by naked mole-rat-like histone H1.0 proteins may confer significant resistance to both senescence and age-related cancer to otherwise senescence-prone and/or cancer-susceptible multicellular species, including humans.

## 1. INTRODUCTION

### 1.1. Background

The rodent species known as naked mole-rat *(Heterocephalus glaber*) has been reported to be very long-lived^1^. Moreover, the species is an exception to the age-dependent component of the Gompertz-Makeham empirical law of mortality, which positively correlates mortality rate with age after adulthood^2^. In other words, this species is not only very long-lived but also appears not to undergo any significant senescence process after reaching adulthood. The naked mole-rat also displays an almost negligible cancer incidence^3^—the only unambiguously diagnosed case in this species corresponds to a gastric neuroendocrine carcinoma^4^.

Research on cancer resistance in the naked mole-rat has alraedy identified a mediator in its fibroblastic tissue^5^, along with cancer-resistance specific (i) gene products^6,7^ and (ii) oncogen disruptions^7^. However, the fundamental reason why the naked mole-rat does not undergo senescence nor display significant cancer incidence remains unclear, let alone how to artificially reproduce such remarkable traits in senescence-prone and/or cancer-susceptible species such as ours. This paper intends to address both questions.

The recently proposed hologenic theory of individuated multicellularity^8^ and a theory of senescence derived from it^9^ may shed some light onto the underpinnings of this senescence and cancer resistance. Both theories are grounded in considerations of thermodynamic constraints on histone post-translational modifications. Based on these two theoretical descriptions, I offer an experimentally falsifiable theory that may explain the naked mole-rat’s remarkable negligible-senescence and cancer-resistance traits in terms of chromatin dynamics. The plausibility of this theory is here supported by three unambiguous proofs of concept. If it resists falsification attempts consistently, the theory will be important for a better understanding of senescence and cancer processes in the naked mole-rat and other individuated multicellular species. The theory may also inform biotechnological research and could ultimately be important for potential therapeutic or even prophylactic options for senescence-related health conditions in biomedicine.

### 1.2. Theoretical considerations

Histone post-translational modifications (hPTMs), observable in nucleosome core particles (NCPs) nearby each transcription start site (TSS), can be understood as a physical medium with finite capacity to convey biologically meaningful information content. For instance, a fraction of this information content has been used to robustly predict transcript abundance levels from TSS-adjacent hPTM levels^10^. According to the hologenic theory of individuated multicellularity, two mutually independent constraints on hPTMs in each NCP respectively convey two types of critical yet mutually unrelated information contents necessary for transcriptional regulation at the multicellular-individual level^8^. Here, constraints are understood as local and level-of-scale specific thermodynamic boundary conditions.

The first information content is hologenic content, which is conveyed in constraints that depend on interactions between NCP histones and other proteins^8^; this content regulates (i.e., constrains) transcription, making it accurate for the needs of the multicellular individual (Fig. 1a, blue). The second is epigenetic information content, which is conveyed in constraints dependent on interactions between NCP histones and DNA^8^; this content regulates (i.e., constrains) transcription making it precise for said needs (Fig. 1b, orange).

**Fig. 1.**
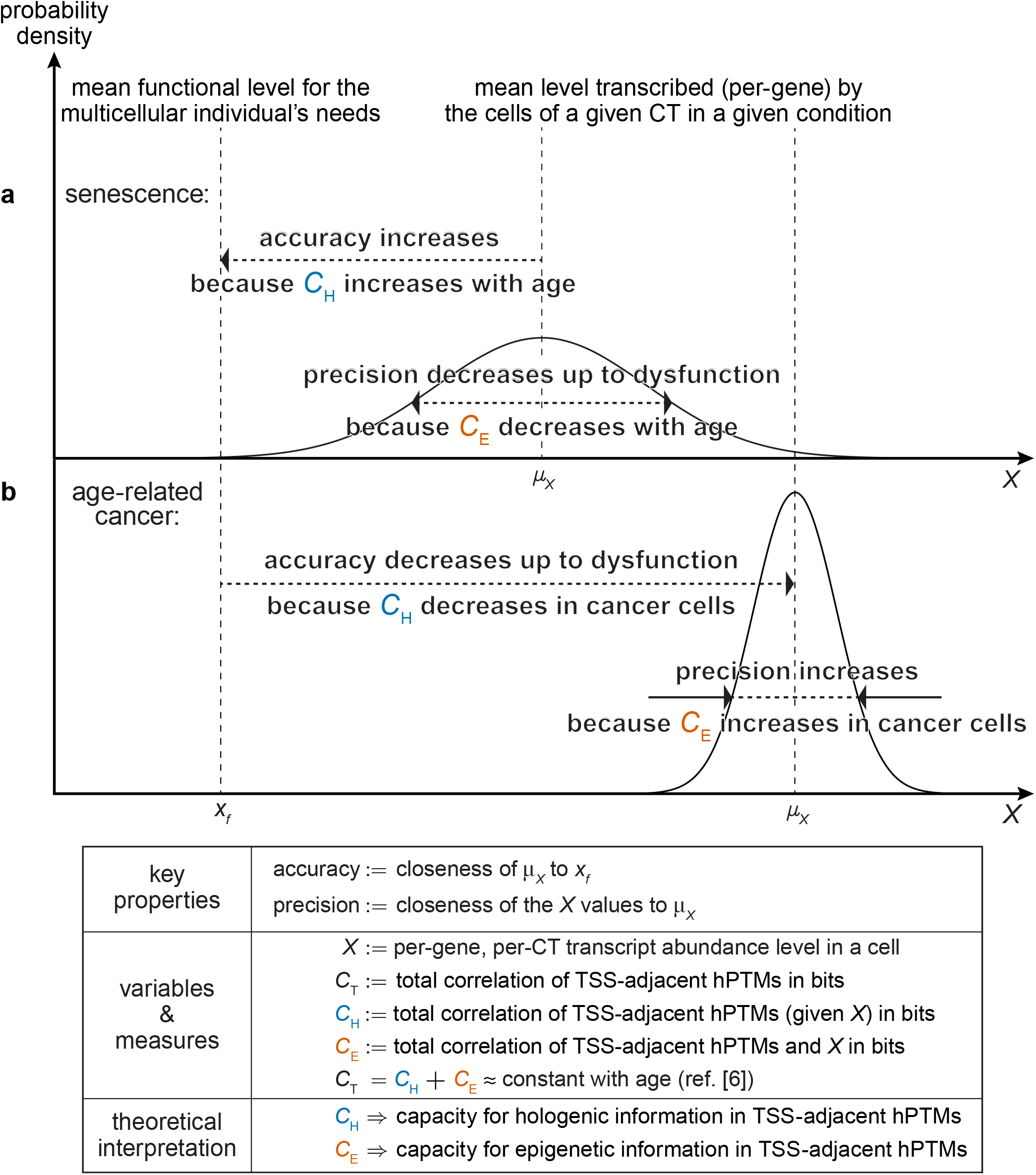
Schematic PDFs representing the theory of senescence as an imbalance of hologenic (blue) and epigenetic (orange) information in hPTMs^9^. (a) Senescence is a developmental byproduct caused by a substantial imbalance in the information conveyed by the constraints on chromatin dynamics—in which transcription becomes increasingly over-regulated after the multicellular individual reaches adulthood, in terms of gaining accuracy at the increasing expense of precision—up to the point of dysfunctionality for the multicellular individual. (b) Age-related cancer is the result of a poorly tuned yet strong enough “pushback” by the multicellular individual against its own senescence process—making transcription dysregulated by gaining precision at the increasing expense of accuracy—up to the point of dysfunctionality for the multicellular individual. Notes: (i) In the absence of senescence and cancer, transcriptional accuracy and transcriptional precision increase/decrease at the expense/gain of each other within ranges that are functional for the multicellular individual’s needs. (ii) Constraints on hPTMs can be quantified in bits using the Shannonian measures known as total correlation *C*(*Y*_1_,...,*Y_n_*) and conditional total correlation *C*(*Y*_1_,..., *Y_n_* |*X* = *x*)^32^, where {*Y*_1_,..., *Y_n_*} are random variables representing TSS-adjacent hPTM levels in the NCP and *X* is a random variable representing the per-gene, per-cell-type transcript abundance level in a cell.

The theory of senescence describes senescence as a developmental byproduct in which hologenic information capacity in hPTMs is increasingly gained in each NCP and throughout development at the expense of capacity for epigenetic information^9^. This uninterrupted process creates a hologenic/epigenetic information imbalance after adulthood in terms of accuracy versus precision in transcriptional regulation.

To illustrate this without loss of generality, let *geneG* be any given gene, let *X* be a random variable representing the *geneG* transcript abundance in a cell within any given cell type (CT), and let *x_f_* be the mean *geneG* transcript abundance level that is functional for the multicellular individual in any given condition. The hologenic/epigenetic information imbalance over-regulates transcription in terms of accuracy (i.e., closeness of the mean *μ_X_* to *x_f_*) gained at the increasing expense of precision (i.e., closeness of the X values to their own mean *μ_X_*) up to the point of dysfunctionality for the multicellular individua^19^ (see schematic probability density function (PDF) in Fig. 1a). Importantly, this predictable age-dependent loss of transcriptional precision has been experimentally observed in different tissues^11–17^.

The theory of senescence also describes age-related cancer to be a result of a poorly tuned yet strong enough “pushback” by the multicellular individual at the chromatin level against its own senescence process, thereby dysregulating transcription. For each TSS within a given cell type, this pushback occurs in terms of precision gained at the increasing expense of accuracy, up to the point of dysfunctionality for the multicellular individua^l9^ (see schematic PDF in Fig. 1b).

### 1.3. Empirical considerations

In the described two theories, histones and their post-translational modifications embody critical constraints on chromatin dynamics for development-related processes, such as senescence and age-related cancer. In particular, constraints on chromatin dynamics in age-related cancer can be understood as a poorly tuned yet strong enough “pushback” against senescence as mentioned previously^9^. One particular histone family, namely the histone H1 (also known as the “linker” histone) family, could explain the particular traits of the naked mole-rat in terms of chromatin dynamics because (i) histone H1 is less evolutionarily conserved and much more variable in size and sequence than the core histone families (H2A, H2B, H3 and H4)^18^ and (ii) histone H1 constrains the accessibility of critical histone-modifying enzymes and chromatin remodeling factors to the NCP^19,20^.

The globular domain of the histone H1 protein folds into a major structural motif known as a “winged” helix-turn-helix (wHTH)^18^, in the form of *α*_1_-*β*_1_-*α*_2_-*α*_3_-*β*_2_-*β*_3_^21^, where the *α_i_* motifs are alpha helices and the *β_j_* motifs are beta sheets. The *α*_3_ motif within the wHTH motif is critical for the histone H1 binding affinity to the NCP^22–25^.

All known variants of the histone H1 are encoded by paralog gene families^26^. In particular, the two variants relevant for the proposed theory—i.e., with “linker” histone function critical for the adult body or soma—are (i) “replication independent” variants (i.e., they are synthesized throughout the cell cycle) and (ii) synthesized chiefly by somatic cells. For all metazoans the common variant is H1.0 (H1 histone family, member 0; also known as H1°, H1(0), H5, H1*δ*, or RI H1) and within metazoans, vertebrates add a second variant H1x (H1 histone family, member X; also known as H1.10^)21,27^. Whereas H1x histones are highly expressed in human neuroendocrine cells and tumors^28^, H1.0 histones accumulate in terminally differentiated mammalian cells^29,30^, accounting for ≈80% of all H1 transcripts^31^.

## 2. PRESENTATION OF THE THEORY

### 2.1. Summary

- The constraints imposed by the naked mole-rat’s histone H1.0 protein on chromatin dynamics in terminally differentiated cells are a necessary condition for its resistance to both senescence and cancer
- Conversely, the constraints imposed by a naked mole-rat-like histone H1.0 protein on chromatin dynamics in terminally differentiated cells are a sufficient condition for a mutant house to display significant resistance to both senescence and cancer

These causal relationships can be explained in that the naked mole-rat histone H1.0 confers particular stabilization to the histone H1.0-NCP binding affinity in terminally differentiated cells. This stabilization is critical to counteract the otherwise increasing hologenic/epigenetic information imbalance after adulthood that has been proposed as the fundamental cause of senescence^9^ (see Fig. 1a and the subsection entitled Detailed theoretical description).

### 2.2. Proofs of concept

The following three proofs of concept are meant to support the plausibility of the theory but in no way should be regarded as evidence for it. The proofs consist of an independent proof I plus its derived proofs II and III, all obtained by surveying publicly available histone H1 protein sequences:

I. If the theory presented here is correct, the differential constraints embodied by the naked mole-rat histone H1.0 protein should be accounted for by at least one differential amino acid residue in the sequence of that histone protein. In other words, at least one amino acid residue in the naked mole-rat histone H1.0 protein should be non-conserved with respect to a highly conserved residue (at the same homologous site) in vertebrate species both closely or distantly related to the naked mole-rat. A multiple alignment of the histone H1.0 reference protein sequences from the naked mole-rat *(Heterocephalus glaber*), Damaraland mole-rat *(Fukomys damarensis*), Norwegian rat *(Rattus norvegicus*), house mouse *(Mus musculus*), human (*Homo sapiens*), Western painted turtle (*Chrysemys picta bellii*), and the African clawed frog *(Xenopus laevis*) reveals that three of such expected sites exist (sites S_1_, S_2_, and S_3_; see Supplemental file 1). Notably, one of these sites (S_2_) is occupied in the naked mole-rat by a non-conserved arginine (R) residue located in a linker-DNA proximal position^25^ within the *α*_3_-motif sequence (relative position #13 in Fig. 2a). This arginine residue is also proximal to a highly-conserved lysine (K) residue (relative position #11 in Fig. 2a). This highly conserved lysine residue is already known to undergo a post-translational acetylation in the fruit fly *Drosophila melanogaster*^33^ (Table S1). Importantly, lysine acetylation neutralizes the otherwise positive electric charge of this residue and also impairs its ability to form hydrogen bonds^34^ thereby decreasing its binding affinity to the negatively-charged DNA^34,35^.
II. The Cnidaria and Mollusca phyla are interesting for the concept-proofing of the theory because some member species—such as the freshwater polyp *(Hydra vulgaris*)^36^ and the ocean quahog *(Arctica islandica*)^37^—have also been found to display negligible senescence. Nematoda (roundworms) is another interesting phylum because adult lifespan varies up to more than 300-fold among some of its known member species^38,39^. Expanding the explanatory scope of the theory to these phyla and given the proof of concept I, it is expected that only in their long-lived member species—such as the cnidarian *Hydra vulgaris,* the mollusc *Arctica islandica,* and the disease-causing nematodes *Onchocerca volvulus* (onchocerciasis^40^), *Brugia malayi* (lymphatic filariasis^41^), and *Loa loa* (loiasis^42^)—differential constraints on chromatin dynamics entail a conserved arginine (R) residue in the respective histone H1.0 *α*_3_-motif sequences. Remarkably, their *α*_3_-motif sequence alignment reveals an arginine residue conserved only in the long-lived species (Fig. 2b). Moreover, this conserved arginine residue is located in a strikingly similar relative position (#12 in Fig. 2b) to that of the non-conserved arginine residue found in the *α*_3_-motif sequence of the naked mole-rat histone H1.0 protein (relative position #13 in Fig. 2a).
III. In terms of a “negative-control” proof of concept, it is worth noting that the naked mole-rat appears to be somewhat susceptible to cancer onset in neuroendocrine cells^4^. This finding implies that the theorized cancer protection conferred to the other cell types by the constraints embodied by the naked mole-rat histone H1.0 protein are either absent or counteracted in neuroendocrine cells. As mentioned previously, the histone H1x protein is highly expressed in human neuroendocrine tumor cells and, importantly, more expressed than the histone H1.0 protein^28^. Given these facts and the proof of concept I, it is then to be expected that the naked mole-rat histone H1x α_3_-motif sequence does not display any non-conserved residues, let alone a non-conserved arginine (R) residue. A multiple sequence alignment of the naked mole-rat’s histone H1x protein against those of other closely or distantly related vertebrates shows just that (Fig. 2c).

**Fig. 2.**
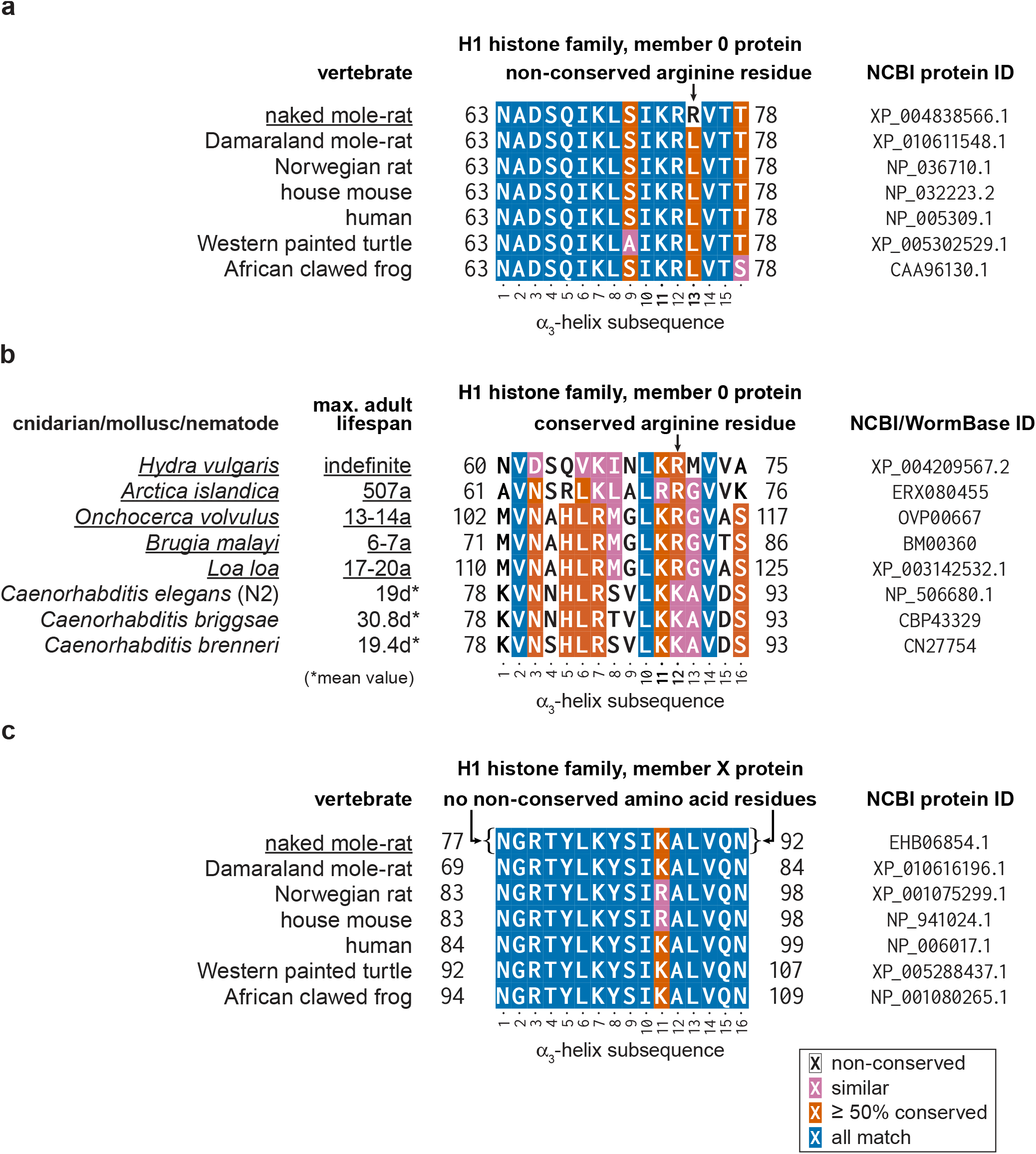
*α*_3_-helix motif subsequence within the multiple sequence alignment of the histone H1.0 and histone H1x proteins. (a) Naked mole-rat’s histone H1.0 *α*_3_-helix subsequence compared to those of other closely or distantly related vertebrate species. (b) Histone H1.0 *α*_3_-helix subsequences of long-lived cnidarian/nematode species compared to those of short-lived nematode species. (c) Naked mole-rat’s histone H1x *α*_3_-helix subsequence compared to those of other closely or distantly related vertebrate species. Multiple sequence alignments were performed using the MAFFT software (v7.419) with its `--globalpair` method option^43^. Identification of the *α*_3_-helix subsequences was done using the curated alignments available in the HistoneDB 2.0 database^21^ as a guide.

### 2.3. Detailed theoretical description

The theory presented here explains the resistance to senescence displayed by the naked mole-rat in the following steps:

- As previous work has shown, H1 histones facilitate chromatin condensation^23^, low histone H1-to-NCP ratios significantly promote chromatin decondensation^44^, and H1 histones also constrain the accessibility to the NCP^20^. In particular, H1 histones constrain hPTM changes in the NCP^44,45^. Thus, with fully functional H1.0 histones in terminally differentiated cells, there is no chromatin decondensation problem, accessibility to the NCP is properly constrained as are hPTM changes in the NCP.
- Given that histone H1.0 proteins accumulate in terminally differentiated cells^26^, the functional-adult scenario changes drastically in senescent multicellular organisms. With age, the highly conserved lysine (K) residue (relative position #11 in Fig. 2a) in the *α*_3_-motif—and/or other post-translationally modifiable residues proximal to it and also to core nucleosomal or linker DNA—lower their positive electric charge as a result of the accumulation of certain post-translational modifications, in particular acetylations^33^ (Table S1 therein) and phosphorylations^46^. Consequently, as the histone H1.0 binding affinity to the negatively-charged DNA in the NCP decreases with age, all the aforementioned constraints progressively dissipate nearby all TSSs. This means all NCPs become more and more accessible, thereby building up hologenic constraints on hPTMs, at the expense of capacity for epigenetic constraints on them—in turn making the dysfunctional over-regulation of transcription (Fig. 1a) a chromatin-scale problem in all terminally differentiated cells.
- In the naked mole-rat, however, the extra arginine (R) residue in the histone H1.0 *α*_3_ motif (relative position #13 in Fig. 2a) creates an additional “reservoir” of positive electric charge in terminally differentiated cells. This arginine-based “reservoir” is possible because arginine is the most basic amino acid residue^47^, displaying a virtually permanent positive charge (i.e., it is always protonated) under physiological conditions^48–50^.
- This “reservoir” of positive electric charge stabilizes histone H1.0-NCP binding affinities when the highly conserved lysine (K) residues (relative position #11 in Fig. 2a)—and/or other amino acid residues proximal to core nucleosomal or linker DNA—accumulate post-translational acetylations or phosphorylations with age (in particular, lysine residues in H1 histones undergo post-translational modifications, including acetylation, as described in different multicellular phyla^33,46,51–54^).
- In other words, histone H1.0-NCP binding affinities near TSSs in the naked mole-rat’s terminally differentiated cells do not decrease significantly with age as is the case for those of senescent multicellular organisms. Such a histone H1.0-NCP binding affinity stabilization is highly dynamic: in general, the mean residency time of H1 histones at any given NCP is less than five minutes^55,56^—much shorter than that of core histones, but in general, longer than that of other chromatin-binding proteins, including transcription factors^57^.
- The post-adulthood naked mole-rat chromatin thus remains under a highly dynamic yet stable constraint equilibrium. In this equilibrium, the respective capacities for hologenic and epigenetic information remain balanced—in particular by keeping hologenic information capacity growth at bay—which in turn maintains both transcriptional accuracy and transcriptional precision within functional ranges (as opposed to what happens in the senescence process; Fig. 1a). The stabilization of chromatin dynamics described here would also explain the observations that (i) naked mole-rat cells display resistance to iPSC (induced pluripotent stem cell) reprogramming^58^ and (ii) even when said reprogramming is successful, the obtained cells are still tumor resistant^7^. In summary, the otherwise increasing hologenic/epigenetic information imbalance that has been proposed to be the fundamental cause of senescence^9^ is corrected by the particular constraints that stabilize the H1.0-NCP binding affinities in the naked mole-rat. Therefore, naked mole-rats display negligible senescence.
- These particular stabilizing constraints are also the reason why naked mole-rats display almost negligible cancer incidence: its histone H1.0-NCP binding affinities are highly stable and there is no significant senescence process that may force this rodent to “push back” against senescence at the chromatin level (see also Theoretical considerations and Fig. 1b). A related description of senescence in evolutionary terms has been presented previously^9^.

## 3. TESTING THE THEORY

To test the theory presented here, I provide the following two experimentally falsifiable predictions:

A. The constraints on chromatin dynamics embodied by a mutant histone H1.0 protein, as specified by the NCBI protein ID XP_004838566.1 with the site-directed amino acid substitution R75L, will negate in the mutant naked mole-rats their otherwise significant resistance to both senescence and cancer.
B. The constraints on chromatin dynamics embodied by a mutant histone H1.0 protein, as specified by the NCBI protein ID NP_032223.2 with the site-directed amino acid substitution L75R, will confer to mutant house mice significant resistance to both senescence and cancer.

## 4. CAN THE SCOPE OF THE THEORY BE EXPANDED TO OTHER SPECIES?

If prediction B holds, it would not be surprising that artificially stabilizing the higher-order constraints on chromatin dynamics in short-lived species other than the house mouse also renders them significantly more resistant to senescence. In other words, having said higher-order constraints embodied by a mutant histone H1.0 (or by a mutant histone H1.0 ortholog) exhibiting an electrostatic DNA binding affinity that remains highly stable throughout adulthood. Such stability is a potentially defining feature of non-senescent or extremely long-lived species from different taxa, as Fig. 3 strongly suggests.

**Fig. 3.**
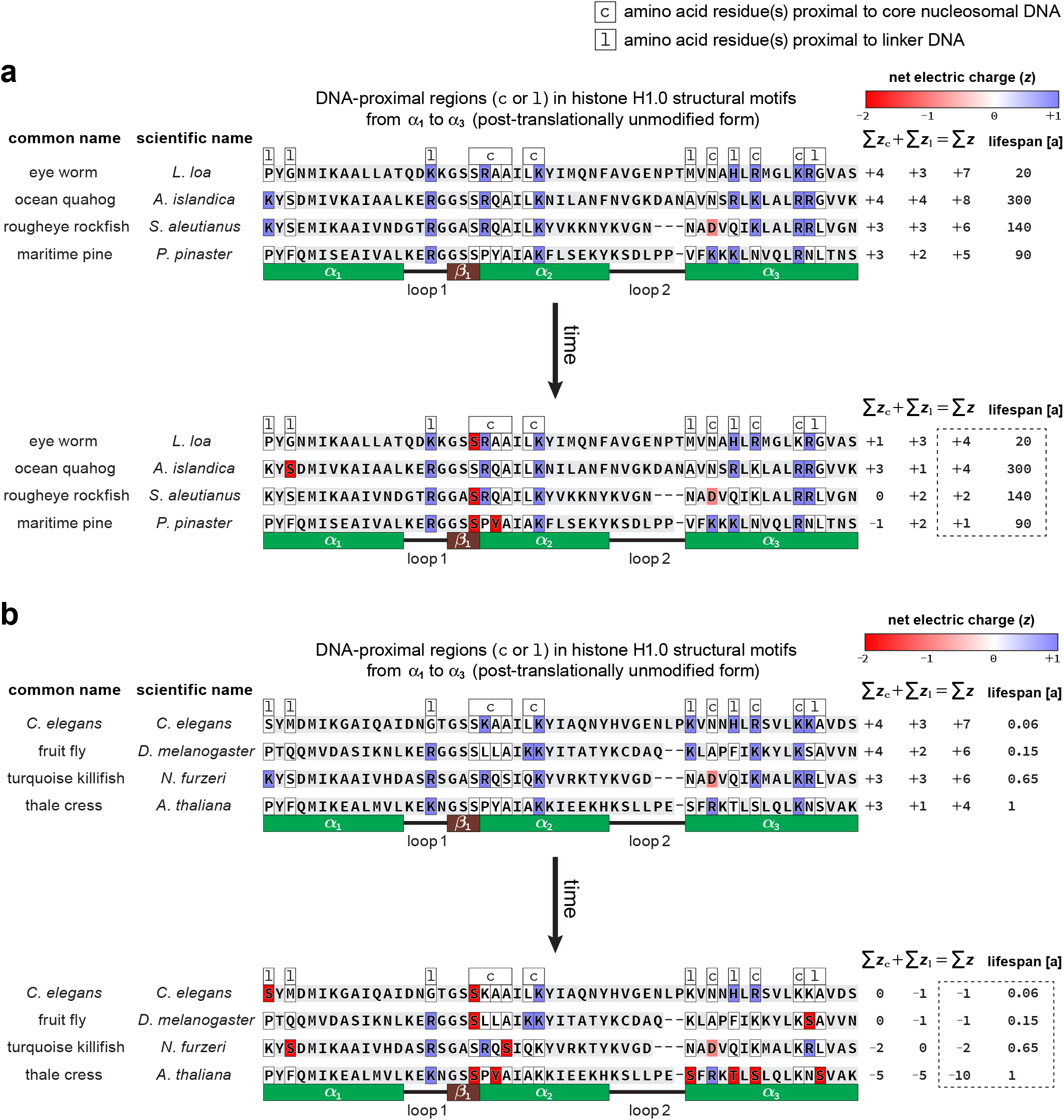
Expanding the scope of the theory to different taxa. (a) In long-lived species, DNA-proximal regions within histone H1.0 protein orthologs accumulate post-translational modifications (in this analysis, Lys acetylations and Ser/Thr/Tyr phosphorylations) throughout adulthood and the associated decrease in electrostatic DNA binding affinity is not substantial, i.e., it does not entail electrostatic repulsion forces on the negatively-charged DNA (b) In short lived species, DNA-proximal regions within histone H1.0 protein orthologs also accumulate post-translational modifications throughout adulthood but the associated decrease in electrostatic DNA binding affinity is substantial, i.e., it does entail electrostatic repulsion forces on the negatively-charged DNA. Metadata and hPTM prediction software used can be found in Supplemental file 2. [Simplifications: Exact, site-specific distances from DNA-proximal amino acid residues to the DNA backbone an allometric relationships were not taken into account. Amino acid residues were assumed to be independent of each other and *z* (electric charge) values were summed on a residue-by-residue basis.]

## 5. IMPLICATIONS OF THE THEORY

The following discussion is made under the assumption that the theory presented here will resist falsification attempts consistently—including the verification of the main predictions A and B provided here.

### 5.1. Basic science implications

The work presented in this article was conducted under two specific assumptions: the constraint-based theoretical descriptions of (i) the evolution of individuated multicellular organisms and their developmental self-regulatory dynamics^8^ and (ii) age-related cancer as the result of a poorly tuned yet strong enough “pushback” against senescence^9^ are both correct.

Given the highly specific nature of the predictions A and B, their verification should make the fundamental approach underpinning the work presented here worth some consideration. Indeed, the generic form of this approach has been around for more than a decade and its great explanatory power is still largely underused. To a large extent, this subsection aims to address this issue.

To date, a number of theorists, myself included, have argued at lengths that phenomena such as abiogenesis^59,60^, individuated multicellularity^8^, senescence and cancer^9^, mental processes^61^, biological complexity^62,63^, and the life phenomenon itself^61,64^ should be fundamentally understood in terms of an emergent, higher-order constraint (i.e., a constraint on constraints or *teleodynamic* constraint) on the release of energy—thereby performing teleodynamic work^61^. Notably, this approach has also been independently framed and discussed in terms of a “closure of constraints”^65–67^.

Higher-order or teleodynamic constraints emerge from and supervene on synergistically coupled lower-order constraints^61^. The supervenience concept—applied here in its simplest form^68^—means there can be no change in a higher-order constraint without a change in some of the lower-order constraints it emerges from. In this sense, the naked mole-rat’s resistance to both senescence and cancer is supervenient on lower-order constraints. These lower-order constraints can in turn be drastically altered with the build-up of constraints embodied by something as small and subtle as a very specific amino acid residue in a very specific motif within a very specific protein encoded by a very specific gene.

Thus, under this view of living systems—grounded on teleodynamics—we should avoid understanding any gene, protein, motif, or amino acid residue as *the* senescence gene, protein, motif, or amino acid residue. Instead, we should understand living systems, and in particular development-related processes, in terms of higher-order constraints. Such constraints are local and level-of-scale specific thermodynamic boundary conditions that are multiply realizable on lower-order molecular dynamics. Although higher-order constraints harness the release of energy into work to preserve themselves, they can be completely dissipated (i.e., death), disrupted (e.g., an insult to a tissue or organ), or subject to a constraint imbalance such as that described for senescence.

### 5.2. Biotechnological and biomedical implications

A direct implication of the theory (*sensu stricto,* if its prediction B holds) is the prospect of modifying the natural constraints on chromatin dynamics—by engineering replication-independent H1 histones and the nucleic acid sequences encoding them—that may confer negligible senescence to species other than the house mouse (via genome editing), including our own (via mRNA delivery). Importantly, stopping senescence and eliminating the incidence of age-related cancer have been proposed to be one and the same technical challenge^9^. Thus, the verification of prediction B may open the door to the development of efficient therapeutic, and even prophylactic, applications for the group of diseases we call cancer.

The theory presented in this paper does not, however, account for lifespan differences among species with identical amino acid residues proximal to core nucleosomal or linker DNA (e.g., among mammals other than the naked mole-rat). In turn, this caveat translates into the question of how to confer resistance to senescence and/or age-related cancer without involving a mutant histone H1.0 protein. Notably, the theory itself may provide clues for achieving just that.

If the α-helical motifs of a wild-type histone H1.0 protein are a target of post-translational modifications as the theory presented here describes, one may consider to inhibit—once the organism has reached its adult form—its phosphorylation. This is because phosphorylation greatly reduces the net electric charge of the serine (S), threonine (T), or tyrosine (Y) residues it modifies. In this context, it is already known that the histone H1 protein is a target of cyclin/CDK (cyclin dependent kinase) phosphorylation^69^. Similarly to how the histone H1.0 was singled out in this work because it is replication-independent, CDK5, CDK7, and CDK9 can be identified as suitable targets for highly selective inhibition because they are known to be cell cycle-independent^70^.

Among these three kinases, CDK9^71^ and CDK_5_^72^ are known to phosphorylate H1 histones. Like the histone H1.0, CDK9 has very low tissue specificity and it is found in higher proportions in terminally differentiated cells^73^ (and references therein). CDK5, on the other hand, has high tissue specificity—it is found in higher proportions in the nervous system^74^ (and references therein).

## 6. A SECONDARY PREDICTION

From these considerations, I offer the following secondary prediction: adults from multicellular, senescence-prone and/or cancer-susceptible species will display resistance to both senescence and age-related cancer when treated periodically (e.g., daily) with a highly selective CDK9 inhibitor in minimal doses, and will display additional resistance to both senescence and age-related cancer in their nervous system (if applicable for the given species) when treated periodically with a highly selective CDK5 inhibitor in minimal doses.

Notwithstanding this secondary prediction is speculative—it has a a plausible rationale but no proof of concept—I chose to present it in this paper. The reason is previous research work has found (i) highly selective CDK9 inhibitors to be potentially useful to treat cancer^75,76^ and (ii) roscovitine, a highly selective CDK5 inhibitor, to have potential for neuroprotection^77^ and cancer treatment^78^. Importantly, the fundamental approach underpinning these previous findings is completely unrelated to the fundamental approach described in this paper. This convergence towards the use of selective CDK9 or CDK5 inhibitors is, I argue, remarkable and makes the testing of this secondary prediction worth serious consideration. It must also be noted, however, that a falsification of this prediction does not imply the falsification of the theory presented here as a whole.

## CONCLUDING REMARKS

Whereas the prospect of using highly selective CDK9 or CDK5 inhibitor drugs to confer resistance to senescence might seem compelling due to its simplicity, a safety issue arises. Namely, CDK9 or CDK5 inhibition will not be histone-H1.0 specific. That is, other targets of phosphorylation in the cell nucleus may not be properly phosphorylated whenever this post-translational modification is indeed necessary. In this context, the use of mutant histone H1.0 protein—via mRNA delivery or genome edition in non-human species and only via mRNA delivery in humans—appears to be a much safer option, however complex this strategy might be in technical terms.

## Supporting information

Complete protein sequence aligment for Fig. 2a

Protein sequence metadata for Fig. 3

## ABBREVIATIONS

H1.0/H1°/H1(0)/H5/H1*δ*/RI H1: H1 histone family, member 0;
hPTMs: histone post-translational modifications;
NCP: nucleosome core particle;
TSS: transcription start site;
CT: cell type;
PDF: probability density function;
wHTH: “winged” helix-turn-helix structural motif;
*α*_3_: third (from N-to C-terminus) alpha-helix motif within the major wHTH motif;
H1x/H.10: H1 histone family, member X;
iPSC: induced pluripotent stem cell;
CDK: cyclin-dependent kinase.

## AVAILABILITY OF SUPPORTING DATA

All sequence data analyzed are available in the NCBI Protein, NCBI SRA, and WormBase public repositories. The protein 3D structure data file analyzed is available in the RCSB Protein Data Bank (PDB ID 5NL0).

## ACKNOWLEDGEMENTS

I wish to thank Angelika H. Hofmann at SciWri Services for valuable comments and edits of this paper into an English I could only dream to write. For reviewing this paper and their valuable comments I am indebted to Amy Brock, Daniela Huerta, Mel Andrews, Inti Pedroso, Mauricio Fernandez, Stuart A. Kauffman, and Terrence W. Deacon.

## REFERENCES

1 R. Buffenstein, The naked mole-rat: a new long-living model for human aging research, Journals Gerontol. Ser. A Biol. Sci. Med. Sci. 60 (11) (2005) 1369–1377. doi:10.1093/gerona/60.11.1369.

2 J. G. Ruby, M. Smith, R. Buffenstein, Naked mole-rat mortality rates defy Gompertzian laws by not increasing with age, Elife 7 (2018) 1–18. doi:10.7554/eLife.31157.

3 R. Buffenstein, Negligible senescence in the longest living rodent, the naked mole-rat: Insights from a successfully aging species, J. Comp. Physiol. B Biochem. Syst. Environ. Physiol. 178 (4) (2008) 439–445. doi:10.1007/s00360-007-0237-5.

4 M. A. Delaney, J. M. Ward, T. F. Walsh, S. K. Chinnadurai, K. Kerns, M. J. Kinsel, P. M. Treuting, Initial case reports of cancer in naked mole-rats *(Heterocephalus glaber*), Vet. Pathol. 53 (3) (2015) 691–696. doi:10.1177/0300985816630796.

5 X. Tian, J. Azpurua, C. Hine, A. Vaidya, M. Myakishev-Rempel, J. Ablaeva, Z. Mao, E. Nevo, V. Gorbunova, A. Seluanov, High-molecular-mass hyaluronan mediates the cancer resistance of the naked mole rat, Nature 499 (7458) (2013) 346–349. arXiv:NIHMS150003, doi:10.1038/nature12234. URL http://dx.doi.org/10.1038/nature12234

6 X. Tian, J. Azpurua, Z. Ke, A. Augereau, Z. D. Zhang, J. Vijg, V. N. Gladyshev, V. Gorbunova, A. Seluanov, INK4 locus of the tumor-resistant rodent, the naked mole rat, expresses a functional p15/p16 hybrid isoform, Proc. Natl. Acad. Sci. U. S. A. 112 (4) (2015) 1053–1058. doi:10.1073/pnas.1418203112.

7 S. Miyawaki, Y. Kawamura, Y. Oiwa, A. Shimizu, T. Hachiya, H. Bono, I. Koya, Y. Okada, T. Kimura, Y. Tsuchiya, S. Suzuki, N. Onishi, N. Kuzumaki, Y. Matsuzaki, M. Narita, E. Ikeda, K. Okanoya, K. I. Seino, H. Saya, H. Okano, K. Miura, Tumour resistance in induced pluripotent stem cells derived from naked mole-rats, Nat. Commun. 7 (May) (2016) 1–9. doi:10.1038/ncomms11471.

8 F. A. Veloso, On the developmental self-regulatory dynamics and evolution of individuated multicellular organisms, J. Theor. Biol. 417 (2017) 84–99. doi:10.1016/j.jtbi.2016.12.025.

9 F. A. Veloso, A constraints-based theory of senescence: imbalance of epigenetic and non-epigenetic information in histone crosstalk, bioRxiv (2018) doi:10.1101/310300.

10 V. Kumar, M. Muratani, N. A. Rayan, P. Kraus, T. Lufkin, H. H. Ng, S. Prabhakar, Uniform, optimal signal processing of mapped deep-sequencing data, Nat. Biotechnol. 31 (7) (2013) 615–622. doi:10.1038/nbt.2596.

11 M. C. Salzer, A. Lafzi, A. Berenguer-Llergo, C. Youssif, A. Castellanos, G. Solanas, F. O. Peixoto, C. Stephan-Otto Attolini, N. Prats, M. Aguilera, J. Martín-Caballero, H. Heyn, S. A. Benitah, Identity Noise and Adipogenic Traits Characterize Dermal Fibroblast Aging, Cell 175 (6) (2018) 1575–1590.e22. doi:10.1016/j.cell.2018.10.012.

12 R. Bahar, C. H. Hartmann, K. A. Rodriguez, A. D. Denny, R. A. Busuttil, M. E. Dolle, R. B. Calder, G. B. Chisholm, B. H. Pollock, C. A. Klein, J. Vijg, Increased cell-to-cell variation in gene expression in ageing mouse heart, Nature 441 (7096) (2006) 1011–1014. doi:10.1038/nature04844.

13 I. Angelidis, L. M. Simon, I. E. Fernandez, M. Strunz, C. H. Mayr, F. R. Greiffo, G. Tsitsiridis, M. Ansari, E. Graf, T. M. Strom, M. Nagendran, T. Desai, O. Eickelberg, M. Mann, F. J. Theis, H. B. Schiller, An atlas of the aging lung mapped by single cell transcriptomics and deep tissue proteomics, Nat. Commun. 10 (1) (2019) 1–17. doi:10.1038/s41467-019-08831-9. URL http://dx.doi.org/10.1038/s41467-019-08831-9

14 C. Nikopoulou, S. Parekh, P. Tessarz, Ageing and sources of transcriptional heterogeneity, Biol. Chem. 400 (7) (2019) 867–878. doi:10.1515/hsz-2018-0449.

15 C. P. Martinez-Jimenez, N. Eling, H.-C. Chen, C. A. Vallejos, A. A. Kolodziejczyk, F. Connor, L. Stojic, T. F. Rayner, M. J. T. Stubbington, S. A. Teichmann, Others, Aging increases cell-to-cell transcriptional variability upon immune stimulation, Science 355 (6332) (2017) 1433–1436. doi:10.1126/science.aah4115.

16 M. Enge, H. E. Arda, M. Mignardi, J. Beausang, R. Bottino, S. K. Kim, S. R. Quake, Single-Cell Analysis of Human Pancreas Reveals Transcriptional Signatures of Aging and Somatic Mutation Patterns, Cell 171 (2) (2017) 321–330.e14. doi:10.1016/j.cell.2017.09.004.

17 A. Swisa, K. H. Kaestner, Y. Dor, Transcriptional Noise and Somatic Mutations in the Aging Pancreas, Cell Metab. 26 (6) (2017) 809–811. doi:10.1016/j.cmet.2017.11.009. URL https://doi.org/10.1016/j.cmet.2017.11.009

18 H. E. Kasinsky, J. D. Lewis, J. B. Dacks, J. Ausio, Origin of H1 linker histones, FASEB J. 15 (1) (2001) 34–42. doi: 10.1096/fj.00-0237rev.

19 N. Happel, D. Doenecke, Histone H1 and its isoforms: contribution to chromatin structure and function, Gene 431 (1-2) (2009) 1–12. doi:10.1016/j.gene.2008.11.003.

20 F. Song, P. Chen, D. Sun, M. Wang, L. Dong, D. Liang, R.-M. Xu, P. Zhu, G. Li, Cryo-EM study of the chromatin fiber reveals a double helix twisted by tetranucleosomal units, Science 344 (6182) (2014) 376–380. doi:10.1126/science.1251413.

21 E. J. Draizen, A. K. Shaytan, L. Mariño-Ramírez, P. B. Talbert, D. Landsman, A. R. Panchenko, HistoneDB 2.0: a histone database with variants—an integrated resource to explore histones and their variants, Database 2016 (2016) 1–10. doi: 10.1093/database/baw014.

22 R.-M. Mannermaa, J. Oikarinen, A DNA-binding homeodomain in histone H1, Biochem. Biophys. Res. Commun. 168 (1) (1990) 254–260. doi:10.1016/0006-291X(90)91701-S.

23 D. T. Brown, T. Izard, T. Misteli, Mapping the interaction surface of linker histone H10 with the nucleosome of native chromatin in vivo, Nat. Struct. Mol. Biol. 13 (3) (2006) 250–255. doi:10.1038/nsmb1050.

24 B. R. Zhou, J. Jiang, H. Feng, R. Ghirlando, T. S. Xiao, Y. Bai, Structural mechanisms of nucleosome recognition by linker histones, Mol. Cell 59 (4) (2015) 628–638. doi:10.1016/j.molcel.2015.06.025.

25 J. Bednar, I. Garcia-Saez, R. Boopathi, A. R. Cutter, G. Papai, A. Reymer, S. H. Syed, I. N. Lone, O. Tonchev, C. Crucifix, H. Menoni, C. Papin, D. A. Skoufias, H. Kurumizaka, R. Lavery, A. Hamiche, J. J. Hayes, P. Schultz, D. Angelov, C. Petosa, S. Dimitrov, Structure and dynamics of a 197 bp nucleosome in complex with linker histone H1, Mol. Cell 66 (3) (2017) 384–397.e8. doi:10.1016/j.molcel.2017.04.012.

26 L. Millán-Ariño, A. Izquierdo-Bouldstridge, A. Jordan, Specificities and genomic distribution of somatic mammalian histone H1 subtypes, Biochim. Biophys. Acta – Gene Regul. Mech. 1859 (3) (2016) 510–519. doi:10.1016/j.bbagrm.2015.10.013.

27 P. B. Talbert, K. Ahmad, G. Almouzni, J. Ausió, F. Berger, P. L. Bhalla, W. M. Bonner, W. Z. Cande, B. P. Chadwick, S. W. L. Chan, Others, A unified phylogeny-based nomenclature for histone variants, Epigenetics & Chromatin 5 (1) (2012) 7. doi:10.1186/1756-8935-5-7.

28 J. Warneboldt, F. Haller, O. Horstmann, B. C. Danner, L. Füzesi, D. Doenecke, N. Happel, Histone H1x is highly expressed in human neuroendocrine cells and tumours, BMC Cancer 8 (388) (2008) 1–9. doi:10.1186/1471-2407-8-388.

29 W. Helliger, H. Lindner, O. Grübl-Knosp, B. Puschendorf, Alteration in proportions of histone H1 variants during the differentiation of murine erythroleukaemic cells, Biochem. J. 288 (3) (1992) 747–751. doi:10.1042/bj2880747.

30 J. Zlatanova, D. Doenecke, Histone H1 zero: a major player in cell differentiation?, FASEB J. 8 (15) (1994) 1260–1268. doi: 10.1096/fasebj.8.15.8001738.

31 J.-M. Terme, B. Sesé, L. Millán-Ariño, R. Mayor, J. C. I. Belmonte, M. J. Barrero, A. Jordan, Histone H1 variants are differentially expressed and incorporated into chromatin during differentiation and reprogramming to pluripotency, J. Biol. Chem. 286 (41) (2011) 35347–35357. doi:10.1074/jbc.M111.281923.

32 S. Watanabe, Information theoretical analysis of multivariate correlation, IBM J. Res. Dev. 4 (1) (1960) 66–82. doi:10.1147/rd.41.0066.

33 B. T. Weinert, S. A. Wagner, H. Horn, P. Henriksen, W. R. Liu, J. V. Olsen, L. J. Jensen, C. Choudhary, Proteome-wide mapping of the *Drosophila* acetylome demonstrates a high degree of conservation of lysine acetylation, Sci. Signal. 4 (183) (2011) 1–12. doi:10.1126/scisignal.2001902.

34 X. J. Yang, E. Seto, Lysine acetylation: codified crosstalk with other posttranslational modifications, Mol. Cell 31 (4) (2008) 449–461. doi:10.1016/j.molcel.2008.07.002.

35 J. Ren, Y. Sang, Y. Tan, J. Tao, J. Ni, S. Liu, X. Fan, W. Zhao, J. Lu, W. Wu, Y. F. Yao, Acetylation of lysine 201 inhibits the DNA-binding ability of PhoP to regulate *Salmonella* virulence, PLoS Pathog. 12 (3) (2016) e1005458. doi:10.1371/journal.ppat.1005458.

36 R. Schaible, A. Scheuerlein, M. J. Dańko, J. Gampe, D. E. Martinez, J. W. Vaupel, Constant mortality and fertility over age in *Hydra*, Proc. Natl. Acad. Sci. 112 (51) (2015) 201521002. doi:10.1073/pnas.1521002112.

37 D. Munro, P. U. Blier, The extreme longevity of Arctica islandica is associated with increased peroxidation resistance in mitochondrial membranes, Aging Cell 11 (5) (2012) 845–855. doi:10.1111/j.1474-9726.2012.00847.x.

38 J. Joyner-Matos, A. Upadhyay, M. P. Salomon, V. Grigaltchik, C. F. Baer, Genetic (co)variation for life span in rhabditid nematodes: role of mutation, selection, and history, J. Gerontol. A Biol. Sci. Med. Sci. 64 (11) (2009) 1134–1145. doi:10.1093/gerona/glp112.

39 D. Gems, Longevity and ageing in parasitic and free-living nematodes, Biogerontology 1 (4) (2000) 289–307. doi:10.1023/A:1026546719091.

40 A. Crump, C. M. Morel, S. Omura, The onchocerciasis chronicle: from the beginning to the end?, Trends Parasitol. 28 (7) (2012) 280–288. doi:10.1016/j.pt.2012.04.005.

41 P. J. Lammie, G. Weil, R. Noordin, P. Kaliraj, C. Steel, D. Goodman, V. B. Lakshmikanthan, E. Ottesen, Recombinant antigen-based antibody assays for the diagnosis and surveillance of lymphatic filariasis—a multicenter trial, Filaria J. 3 (1) (2004) 9. doi:10.1186/1475-2883-3-9.

42 J. J. Padgett, K. H. Jacobsen, Loiasis: African eye worm, Trans. R. Soc. Trop. Med. Hyg. 102 (10) (2008) 983–989. doi:10.1016/j.trstmh.2008.03.022.

43 K. Katoh, H. Toh, Recent developments in the MAFFT multiple sequence alignment program, Brief. Bioinform. 9 (4) (2008) 286–298. doi:10.1093/bib/bbn013.

44 Y. Fan, T. Nikitina, J. Zhao, T. J. Fleury, R. Bhattacharyya, E. E. Bouhassira, A. Stein, C. L. Woodcock, A. I. Skoultchi, Histone H1 depletion in mammals alters global chromatin structure but causes specific changes in gene regulation, Cell 123 (7) (2005) 1199–1212. doi:10.1016/j.cell.2005.10.028.

45 J. E. Herrera, Y. Nakatani, R. L. Schiltz, K. L. West, M. Bustin, Histone H1 is a specific repressor of core histone acetylation in chromatin, Mol. Cell. Biol. 20 (2) (2002) 523–529. doi:10.1128/mcb.20.2.523-529.2000.

46 J. R. Wiśniewski, A. Zougman, S. Kruger, M. Mann, Mass spectrometric mapping of linker histone H1 variants reveals multiple acetylations, methylations, and phosphorylation as well as differences between cell culture and tissue, Mol. Cell. Proteomics 6 (1) (2007) 72–87. doi:10.1074/mcp.M600255-MCP200.

47 C. A. Fitch, M. Okon, G. Platzer, B. Garcia-Moreno E., L. P. McIntosh, Arginine: its pKa value revisited, Protein Sci. 24 (5) (2015) 752–761. doi:10.1002/pro.2647.

48 B. Li, I. Vorobyov, A. D. MacKerell, T. W. Allen, Is arginine charged in a membrane?, Biophys. J. 94 (2) (2008) L11–L13. doi:10.1529/biophysj.107.121566.

49 J. Yoo, Q. Cui, Does arginine remain protonated in the lipid membrane? Insights from microscopic pKa calculations, Biophys. J. 94 (8) (2008) L61–L63. doi:10.1529/biophysj.107.122945.

50 M. J. Harms, J. L. Schlessman, G. R. Sue, Others, Arginine residues at internal positions in a protein are always charged, Proc. Natl. Acad. Sci. 108 (47) (2011) 18954–18959. doi:10.1073/pnas.1104808108.

51 C. Bonet-Costa, M. Vilaseca, C. Diema, O. Vujatovic, A. Vaquero, N. Omeñaca, L. Castejón, J. Bernués, E. Giralt, F. Azorín, Combined bottom-up and top-down mass spectrometry analyses of the pattern of post-translational modifications of *Drosophila melanogaster* linker histone H1, J. Proteomics 75 (13) (2012) 4124–4138. doi:10.1016/j.jprot.2012.05.034.

52 A. Izzo, R. Schneider, The role of linker histone H1 modifications in the regulation of gene expression and chromatin dynamics, Biochim. Biophys. Acta – Gene Regul. Mech. 1859 (3) (2016) 486–495. doi:10.1016/j.bbagrm.2015.09.003.

53 B. A. Garcia, S. A. Busby, C. M. Barber, J. Shabanowitz, C. D. Allis, D. F. Hunt, Characterization of phosphorylation sites on histone H1 isoforms by tandem mass spectrometry, J. Proteome Res. 3 (6) (2004) 1219–1227. doi:10.1021/pr0498887.

54 M. Kotliński, K. Rutowicz, L. Knizewski, A. Palusiński, J. Oledzki, A. Fogtman, T. Rubel, M. Koblowska, M. Dadlez, K. Ginalski, A. Jerzmanowski, Histone H1 variants in Arabidopsis are subject to numerous post-translational modifications, both conserved and previously unknown in histones, suggesting complex functions of H1 in plants, PLoS One 11 (1) (2016) 1–19. doi:10.1371/journal.pone.0147908.

55 T. Misteli, A. Gunjan, R. Hock, M. Bustin, D. T. Brown, Dynamic binding of histone H1 to chromatin in living cells, Nature 408 (6814) (2000) 877–881. doi:10.1038/35048610.

56 M. A. Lever, J. P. Th’ng, X. Sun, M. J. Hendzel, Rapid exchange of histone H1.1 on chromatin in living human cells, Nature 408 (6814) (2000) 873–876. doi:10.1038/35048603.

57 T. W. Flanagan, D. T. Brown, Molecular dynamics of histone H1, Biochim. Biophys. Acta – Gene Regul. Mech. 1859 (3) (2016) 468–475. doi:10.1016/j.bbagrm.2015.10.005.

58 L. Tan, Z. Ke, G. Tombline, N. Macoretta, K. Hayes, X. Tian, R. Lv, J. Ablaeva, M. Gilbert, N. V. Bhanu, Z. F. Yuan, B. A. Garcia, Y. G. Shi, Y. Shi, A. Seluanov, V. Gorbunova, Naked mole rat cells have a stable epigenome that resists iPSC reprogramming, Stem Cell Reports 9 (5) (2017) 1721–1734. doi:10.1016/j.stemcr.2017.10.001.

59 T. W. Deacon, Reciprocal linkage between self-organizing processes is sufficient for self-reproduction and evolvability, Biol. Theory 1 (2) (2006) 136–149. doi:10.1162/biot.2006.1.2.136.

60 T. W. Deacon, A. Srivastava, J. A. Bacigalupi, The transition from constraint to regulation at the origin of life, Front. Biosci. 19 (2014) 945–957. doi:10.2741/4259.

61 T. W. Deacon, Incomplete Nature: How Mind Emerged from Matter, W.W. Norton & Company, New York, 2011.

62 S. Koutroufinis, Teleodynamics: A neo-naturalistic conception of organismic teleology, in: B. G. Henning, A. C. Scarfe (Eds.), Beyond Mech. Putt. Life Back Into Biol., Lexington Books, 2013, pp. 312–345.

63 T. Deacon, S. Koutroufinis, Complexity and dynamical depth, Information 5 (3) (2014) 404–423. doi:10.3390/info5030404.

64 T. Deacon, T. Cashman, Teleology versus mechanism in biology: beyond self-organization, in: B. G. Henning, A. C. Scarfe (Eds.), Beyond Mech. Putt. Life Back Into Biol., Lexington Books, 2013, pp. 290–311.

65 M. Montévil, M. Mossio, Biological organisation as closure of constraints, J. Theor. Biol. 372 (2015) 179–191. doi:10.1016/j.jtbi.2015.02.029.

66 M. Mossio, L. Bich, What makes biological organisation teleological?, Synthese 194 (4) (2017) 1089–1114. doi:10.1007/s11229-014-0594-z.

67 S. A. Kauffman, A World Beyond Physics: The Emergence and Evolution of Life, Oxford University Press, New York, USA, 2019.

68 B. McLaughlin, K. Bennett, Supervenience, in: E. N. Zalta (Ed.), Stanford Encycl. Philos., winter 201 Edition, Metaphysics Research Lab, Stanford University, 2018. URL https://plato.stanford.edu/archives/win2018/entries/supervenience

69 A. Contreras, T. K. Hale, D. L. Stenoien, J. M. Rosen, M. A. Mancini, R. E. Herrera, The Dynamic Mobility of Histone H1 Is Regulated by Cyclin/CDK Phosphorylation, Mol. Cell. Biol. 23 (23) (2003) 8626–8636. doi:10.1128/mcb.23.23.8626-8636.2003.

70 K. Wang, P. Hampson, J. Hazeldine, V. Krystof, M. Strnad, P. Pechan, J. M. Lord, Cyclin-dependent kinase 9 activity regulates neutrophil spontaneous apoptosis, PLoS One 7 (1) (2012) 3–8. doi:10.1371/journal.pone.0030128.

71 S. K. O’Brien, H. Cao, R. Nathans, A. Ali, T. M. Rana, P-TEFb kinase complex phosphorylates histone H1 to regulate expression of cellular and HIV-1 genes, J. Biol. Chem. 285 (39) (2010) 29713–29720. doi:10.1074/jbc.M110.125997. URL http://dx.doi.org/10.1074/jbc.M110.125997

72 D. Tang, J. Yeung, K. Y. Lee, M. Matsushita, H. Matsui, K. Tomizawa, O. Hatase, J. H. Wang, An isoform of the neuronal cyclin-dependent kinase 5 (Cdk5) activator, J. Biol. Chem. 270 (45) (1995) 26897–26903. doi:10.1074/jbc.270.45.26897. URL http://dx.doi.org/10.1074/jbc.270.45.26897

73 G. De Falco, A. Giordano, CDK9: From basal transcription to cancer and AIDS, Cancer Biol. Ther. 1 (4) (2002) 342–347. doi:10.4161/cbt.1.4.6113.

74 R. Dhavan, L. H. Tsai, A decade of CDK5, Nat. Rev. Mol. Cell Biol. 2 (10) (2001) 749–759. doi:10.1038/35096019.

75 C. Alcon, A. Manzano-Munoz, J. Montero, A new CDK9 inhibitor on the block to treat hematologic malignancies, Clin. Cancer Res. 26 (4) (2020) 761–763. doi:10.1158/1078-0432CCR-19-3670.

76 X. Wang, C. Yu, C. Wang, Y. Ma, T. Wang, Y. Li, Z. Huang, M. Zhou, P. Sun, J. Zheng, S. Yang, Y. Fan, R. Xiang, Novel cyclin-dependent kinase 9 (CDK9) inhibitor with suppression of cancer stemness activity against non-small-cell lung cancer, Eur. J. Med. Chem. 181 (2019) 111535. doi: https://doi.org/10.1016/j.ejmech.2019.07.038. URL https://www.sciencedirect.com/science/article/pii/S0223523419306592

77 B. Menn, S. Bach, T. L. Blevins, M. Campbell, L. Meijer, S. Timsit, Delayed treatment with systemic (s)-roscovitine provides neuroprotection and inhibits in vivo CDK5 activity increase in animal stroke models, PLoS One 5 (8). doi: 10.1371/journal.pone.0012117.

78 J. Cicenas, K. Kalyan, A. Sorokinas, E. Stankunas, J. Levy, I. Meskinyte, V. Stankevicius, A. Kaupinis, M. Valius, Roscovitine in cancer and other diseases, Ann. Transl. Med. 3 (10) (2015) 1–12. doi:10.3978/j.issn.2305-5839.2015.03.61.

