## Supplementary material for "How the naked mole-rat resists senescence: a constraints-based theory": Complete protein sequence aligment for Fig. 2a

### Multiple protein sequence alignment
### H1.0 (H1 histone family, member 0) proteins in Vertebrata

alignment reproducible with: `mafft --globalpair --maxiterate 5000 sequences.txt > alignment.fa`

site $S_1$ (position 38)

```
naked mole-rat          MTENSTSTPAAKPKRAKASKKSTDHPKYSDMIVAAIQTEKNRAGSSRQSIQKYIKSHYKM   60
Damaraland mole-rat     MTENTTSAPAAKPKRAKASKKSIDHPKYSDMIVAAIQAEKNRAGSSRQSIQKYIKSHYKV   60
Norwegian rat           MTENSTSTPAAKPKRAKAAKKSTDHPKYSDMIVAAIQAEKNRAGSSRQSIQKYIKSHYKV   60
house mouse             MTENSTSAPAAKPKRAKASKKSTDHPKYSDMIVAAIQAEKNRAGSSRQSIQKYIKSHYKV   60
human                   MTENSTSAPAAKPKRAKASKKSTDHPKYSDMIVAAIQAEKNRAGSSRQSIQKYIKSHYKV   60
Western painted turtle  MTENSASAPAAKPKRAKAAKKSTDHPKYSDMIVAAIQAEKSRAGSSRQSIQKYIKSHYKV   60
African clawed frog     MAENSAATPAAKPKRSKALKKSTDHPKYSDMILAAVQAEKSRSGSSRQSIQKYIKNHYKV   60
```

site $S_2$ (position 75)

```
naked mole-rat          GENADSQIKLSIKRRVTTGVLKQTKGVGASGSFRLAKSDEPKRAVAFKKTKKEVKKVATP  120
Damaraland mole-rat     GENADSQIKLSIKRLVTTGVLKQTKGVGASGSFRLAKSDEPKRAVAFKKTKKEVKKVATP  120
Norwegian rat           GENADSQIKLSIKRLVTTGVLKQTKGVGASGSFRLAKGDEPKRSVAFKKTKKEVKKVATP  120
house mouse             GENADSQIKLSIKRLVTTGVLKQTKGVGASGSFRLAKGDEPKRSVAFKKTKKEVKKVATP  120
human                   GENADSQIKLSIKRLVTTGVLKQTKGVGASGSFRLAKSDEPKKSVAFKKTKKEIKKVATP  120
Western painted turtle  GENADSQIKLAIKRLVTTGVLKQTKGVGASGSFRLAKGDEPKKPPPAKKAKKEVKKATTP  120
African clawed frog     GENADSQIKLSIKRLVTSGALKQTKGVGASGSFRLAKADEGKKPA..KKPKKEIKKAVSP  118
```

$\alpha_3$-helix motif (bracket under positions 64–78)

```
naked mole-rat          KKAAKPKKAA.SKAPSKKPKAT..PVKK.AKKKPAAMPKKAKKPKVVKAKPVKASKPKKA  176
Damaraland mole-rat     KKAAKPKKAA.SKAPSKKPKAT..PVKK.AKKKPAATPKKAKKPKVVKAKPVKASKPKKA  176
Norwegian rat           KKAAKPKKAA.SKAPSKKPKAT..PVKK.AKKKPAATPKKAKKPKIVKVKPVKASKPKKA  176
house mouse             KKAAKPKKAA.SKAPSKKPKAT..PVKK.AKKKPAATPKKAKKPKVVKVKPVKASKPKKA  176
human                   KKASKPKKAA.SKAPTKKPKAT..PVKK.AKKKLAATPKKAKKPKTVKAKPVKASKPKKA  176
Western painted turtle  RKAAKPKKAA.TKSPAKKPKAA...AKK.AKKKPAAAPKKAKKPKTVKAKPVKASKPKKA  175
African clawed frog     KKVAKPKKAAKSPAKAKKPKVAEKKVKKVAKKKPAPSPKKAKKTKTVKAKPVRASKVKKA  178
```

site $S_3$ (position 190)

| | | | NCBI protein ID |
|---|---|---|---|
| naked mole-rat | KPVKPKAKSSAKMAGKKK | 194 | XP_004838566.1 |
| Damaraland mole-rat | KPVKPKAKSSARRASKKK | 194 | XP_010611548.1 |
| Norwegian rat | KPVKPKAKSSAKRASKKK | 194 | NP_036710.1 |
| house mouse | KTVKPKAKSSAKRASKKK | 194 | NP_032223.2 |
| human | KPVKPKAKSSAKRAGKKK | 194 | NP_005309.1 |
| Western painted turtle | KPSKPKAKSSAKKSAKKK | 193 | XP_005302529.1 |
| African clawed frog | KPSKPKAKASPKKSGRK. | 195 | CAA96130.1 |

- X non conserved
- X similar
- X ≥ 50% conserved
- X all match
