## Supplementary material for "How the naked mole-rat resists senescence: a constraints-based theory": Protein sequence metadata for Fig. 3

**Sequences for the alignment of histone H1.0 orthologs in different taxa (spanning from helical motif $\alpha_1$ to $\alpha_3$) displayed in Fig. 3**

**From long-lived species:**

**>XP_003142532.1 linker histone H1 and H5 family protein [Loa loa]**
**PYGNMIKAALLATQDKKGSSRAAILKYIMQNFAVGENPTMVNAHLRMGLKRGVAS**
**>SRX5557232 (SRA) [Panulirus argus]**
**KYSSMITDAITALKERGGSSRQAILKYIVATYKVDER--VLNTQCKLALRRGVNS**
**>ERX080455 (SRA) [Arctica islandica]**
**KYSDMIVKAIAALKERGGSSRQAILKNILANFNVGKDANAVNSRLKLALRRGVVK**
**>LHVS01083274.1: 5686-5396 frame-2 [Sebastes aleutianus]**
**KYSEMIKAAIVNDGTRGGASRQAILKYVKKNYKVGN---NADVQIKLALRRLVGN**
**>CAC84682.1 putative histone H1 [Pinus pinaster]**
**PYFQMISEAIVALKERGGSSPYAIAKFLSEKYKSDLPP-VFKKKLNVQLRNLTNS**

**From short-lived species:**

**>NP_506680.1 Histone H1.X [Caenorhabditis elegans]**
**SYMDMIKGAIQAIDNGTGSSKAAILKYIAQNYHVGENLPKVNNHLRSVLKKAVDS**
**>NP_724341.1 histone H1 [Drosophila melanogaster]**
**PTQQMVDASIKNLKERGGSSLLAIKKYITATYKCDAQ--KLAPFIKKYLKSAVVN**
**>XP_015818886.1 PREDICTED: histone H1.0-B [Nothobranchius furzeri]**
**KYSDMIKAAIVHDASRSGASRQSIQKYVRKTYKVGD---NADVQIKMALKRLVAS**
**>NP_179396.1 histone H1-3 [Arabidopsis thaliana]**
**PYFQMIKEALMVLKEKNGSSPYAIAKKIEEKHKSLLPE-SFRKTLSLQLKNSVAK**

■ **α-helix**

■ **β-sheet**

■ **protein loop**

**— alignment gap**

Lys acetylations predicted using GPS-PAIL (version 2.0):
http://pail.biocuckoo.org/online.php

Ser/Thr/Tyr phosphorylations predicted using NetPhos (version 3.1):
https://services.healthtech.dtu.dk/service.php?NetPhos-3.1
